# A dynamic antibacterial T6SS in *Pantoea agglomerans* pv. *betae* delivers a lysozyme-like effector to antagonize competitors

**DOI:** 10.1101/2021.12.03.471080

**Authors:** Andrea Carobbi, Simone Di Nepi, Chaya M. Fridman, Yasmin Dar, Rotem Ben-Yaakov, Isaac Barash, Dor Salomon, Guido Sessa

## Abstract

The type VI secretion system (T6SS) is deployed by numerous Gram-negative bacteria to deliver toxic effectors into neighboring cells. The genome of *Pantoea agglomerans* pv. *betae* (*Pab*) phytopathogenic bacteria contains a gene cluster (T6SS1) predicted to encode a complete T6SS. Using secretion and competition assays, we found that T6SS1 in *Pab* is a functional antibacterial system that allows this pathogen to outcompete rival plant-associated bacteria found in its natural environment. Computational analysis of the T6SS1 gene cluster revealed that antibacterial effector and immunity proteins are encoded within three dynamic genomic islands that harbor arrays of orphan immunity genes or toxin and immunity cassettes. Functional analysis demonstrated that the specialized antibacterial effector VgrG contains a C-terminal catalytically active glucosaminidase domain that is used to degrade prey peptidoglycan. Moreover, we confirmed that a bicistronic unit at the end of the T6SS1 cluster encodes a novel antibacterial T6SS effector and immunity pair. Together, these results demonstrate that *Pab* T6SS1 is an antibacterial system delivering a lysozyme-like effector to eliminate competitors, and indicate that this bacterium contains novel T6SS effectors.

**Significance Statement:** In this work, we describe the identification of a *Pantoea agglomerans* T6SS as an antibacterial determinant used by this phytopathogen to outcompete bacterial rivals. Furthermore, we provide an in-depth analysis of the T6SS gene cluster and the putative effector and immunity genes that comprise it, and we propose explanations for its dynamic evolution and effector diversification in *Pantoea* strains. Lastly, we experimentally validate two predicted effector and immunity pairs, and we demonstrate that one is a potent lysozyme-like toxin.

## INTRODUCTION

The type VI secretion system (T6SS) is a contact-dependent toxin delivery apparatus widespread among Gram-negative bacteria (Bingle *et al*., 2008; Coulthurst, 2019; Wang *et al*., 2019). It is composed of 14 conserved proteins (TssA-M and a PAAR repeat-containing protein) that are encoded by large gene clusters (Boyer *et al*., 2009; Shneider *et al*., 2013). The T6SS delivers toxins, hereafter referred to as effectors, directly into bacterial and eukaryotic cells (Jurėnas and Journet, 2021). Structurally, the T6SS is similar to a contractile phage tail (Nguyen *et al*., 2018; Wang *et al*., 2019). Contraction of an outer sheath tube propels an inner tube, which is decorated with effectors, out of the bacterial cell and into immediately adjacent cells. This secreted tail tube consists of stacked hexameric rings of the Hcp protein (TssD) that are capped with a spike made of a trimer of valine-glycine repeat proteins (VgrG/TssI) bound to a PAAR (Proline-Alanine-Alanine-aRginine) repeat-containing protein that sharpens it (Basler *et al*., 2012, Nguyen *et al*., 2018).

T6SS effectors can be either toxin domains that are fused to the tail tube components Hcp, VgrG, and PAAR, and referred to as specialized effectors, or proteins that are non-covalently bound to the tail tube components and defined as cargo effectors (Jurėnas and Journet, 2021). Depending on the nature of its effector arsenal, the T6SS mediates inter-bacterial competition and/or contribute to virulence against eukaryotes (Jana and Salomon, 2019; Jurėnas and Journet, 2021). T6SS antibacterial effectors may act in the periplasm or in the cytoplasm of the prey cell and carry diverse toxin domains that target vital bacterial cell components, such as nucleic acids, the cell wall and the membrane (e.g., MacIntyre *et al*., 2010; Durand *et al*., 2014; Jiang *et al*., 2016; Ray *et al*., 2017; Jana and Salomon, 2019). T6SS effectors that target eukaryotic cell processes and contribute to bacterial virulence have also been reported (e.g., Pukatzki *et al*., 2006; Durand *et al*., 2014; Jiang *et al*., 2014, 2016; Feria and Valvano, 2020). Interestingly, certain T6SSs have been shown to deliver both antibacterial and virulence effectors, thus implicating them in both bacterial competition and host pathogenicity (Jiang *et al*., 2014; Ray *et al*., 2017). Numerous effector families have been identified to date (e.g., Russell *et al*., 2013; Durand *et al*., 2014; Salomon *et al*., 2014; Dar *et al*., 2018; Jana *et al*., 2019; Hernandez *et al*., 2020) and it is likely that many additional families remain unknown. To avoid T6SS-mediated self-intoxication, bacteria encode immunity proteins, which provide protection against the effectors’ toxicity; the corresponding genes are adjacent to their cognate effector genes in bicistronic units (Hood *et al*., 2010, Whitney *et al*., 2013). Immunity proteins localize to the cellular compartment of action of the cognate effectors and they generally appear to neutralize the effectors rather than protect the target. In fact, in many instances they have been shown to bind to the catalytic domain of effectors and occlude their binding site or lock them in an inactive conformation (Ruhe *et al*., 2020; Jurėnas and Journet, 2021).

T6SSs are also widely distributed in genomes of bacterial plant pathogens such as *Pseudomonas, Xanthomonas, Ralstonia, Agrobacterium*, and *Erwinia* (Boyer *et al*., 2009; Bernal *et al*., 2018; Bayer-Santos *et al*., 2019; Wu *et al*., 2018 and 2021); yet until recently they have received much less attention than their counterparts in animal pathogens (Bernal *et al*., 2018). In many of these organisms, the T6SS does not appear to be primarily used against the plant host, but rather against other members of the microbial community inhabiting the same environmental niche. Several T6SSs of plant pathogenic bacteria were shown to play a role in inter-bacterial competition, such as in *Pseudomonas syringae* (Haapalainen *et al*., 2012), *Acidovorax citrulli* (Levy *et al*., 2018), *Agrobacterium tumefaciens* (Ma *et al*., 2014), and *Pantoea ananatis* (Shyntum *et al*., 2015). However, in most cases the function and biochemical properties of putative secreted effectors were not investigated. Remarkably, non-pathogenic bacteria have been proposed as biocontrol agent to inhibit plant pathogens and prevent diseases (Bernal *et al*., 2018). In support of this possibility, *Pseudomonas putida* KT2440, whose genome includes three T6SS gene clusters, was shown to outcompete plant pathogens *in vitro* and in planta in a T6SS-dependent manner (Bernal *et al*., 2018).

*Pantoea agglomerans* (*Pa*) is an epiphytic and endophytic bacterium widespread in diverse natural and agricultural habitats (Kobayashi and Palumbo, 2000, Lindow and Brandl, 2003). Strains of *Pa* have evolved into tumorigenic pathogens displaying host specificity on various plants by acquiring a pathogenicity plasmid, which contains genes encoding structural components and effectors of the type III secretion system, enzymes of auxin and cytokinins biosynthesis, and multiple insertion sequences (Guo *et al*., 2002; Barash and Manulis-Sasson, 2009). *Pa* strains inflict important losses in beet (*Beta vulgaris*) crops used for human consumption and sugar industry (Burr et al., 1991), and prevent root development of gypsophila (*Gypsophila paniculata*) cuttings employed for flower production (Cooksey, 1986). Two *Pa* pathogenic pathovars were identified: *Pa* pv. *gypsophilae* (*Pag*) and *Pa* pv. *betae* (*Pab*) (Burr *et al*., 1991). *Pag* causes development of galls on gypsophila and triggers a hypersensitive response on beet. *Pab* causes galls on beet as well as on gypsophila, albeit galls on gypsophila caused by *Pab* are morphologically distinguishable from galls caused by *Pag* (Burr *et al*., 1991).

The presence of a T6SS in the genome of *Pa* phytopathogenic strains, its functionality, and possible contribution to interbacterial competition remain unexplored. In addition, the repertoire and biochemical activity of *Pa* T6SS effectors are largely unknown. In this study, we provide evidence that the *Pab* genome contains a T6SS gene cluster, T6SS1, which encodes a complete set of T6SS core-components and accessory proteins, as well as effectors and immunity genes found within dynamic genomic islands. By functional analysis, we demonstrate that T6SS1 is competent in effectors secretion and plays a role in antibacterial competition that may contribute to *Pab* establishment in its environmental niche. We also show that *Pab* VgrG is a specialized T6SS effector, which induces peptidoglycan degradation leading to cell lysis, and is antagonized by its cognate immunity, Vgi. Lastly, we identify a novel antibacterial effector and immunity pair.

## RESULTS

### *The* Pab *genome contains a T6SS gene cluster with diverse effector and immunity genes*

To characterize the T6SS in *Pantoea agglomerans* pv. *betae* (*Pab*), we first identified gene clusters in the *Pab* genome that encode known T6SS components. Analysis of the *Pab* assembled genome (NCBI RefSeq: NZ**_**LXSW01000076.1) revealed that it contains two conserved T6SS gene clusters previously named T6SS1 and T6SS2 (De Maayer *et al*., 2011) (Fig. 1). The T6SS1 cluster encodes a complete T6SS, whereas T6SS2 encodes only a few system components.

**Fig. 1.**
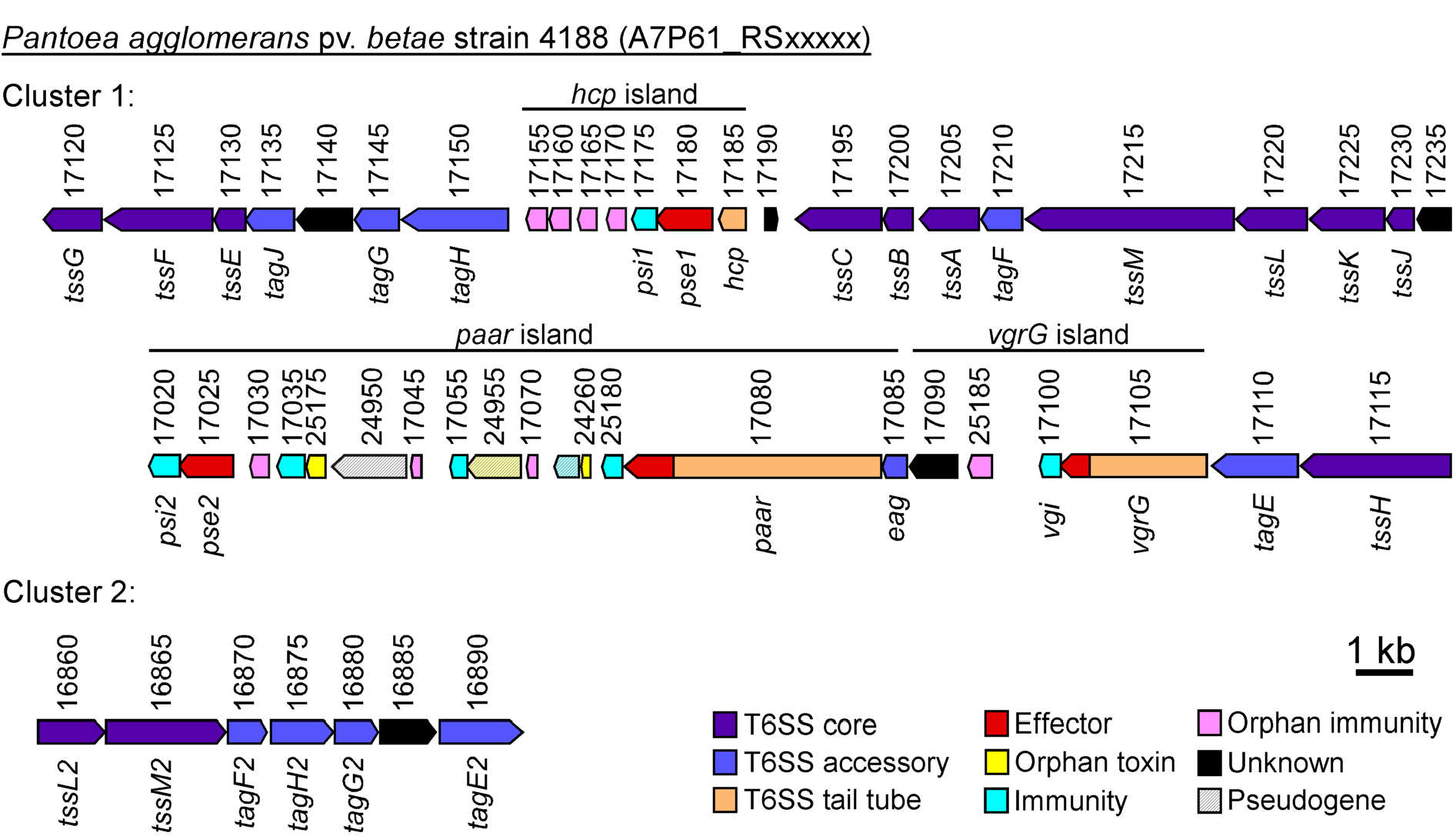
The *Pab* T6SS gene clusters. Schematic representation of T6SS gene clusters present in the *Pab* 4188 genome. Genes are represented by arrows indicating the direction of transcription. Locus tags (A7P61_RSxxxxx) are denoted above. Gene names are shown below. Black lines above denote islands encoding effector and immunity proteins.

#### *hcp* island

T6SS1 includes the previously described *hcp* island between the *hcp* and *tagH* genes (Fig. 1; Supporting Information Fig. S1) (De Maayer *et al*., 2011). The two genes downstream of *hcp* (locus numbers A7P61_RS17180 and A7P61_RS17175) are predicted to encode an antibacterial effector and its cognate immunity, which we termed Pse1 (*<u>P</u>ab* type <u>s</u>ix <u>e</u>ffector <u>1</u>) and Psi1 (*<u>P</u>ab* type <u>s</u>ix immunity <u>1</u>), respectively. Pse1 is similar to pore-forming colicin toxins (97.2% probability of similarity between Pse1 amino acids 102-325 and amino acids 8-197 of the Colicin B C-terminal domain according to HHpred; Gabler *et al*., 2020) and its homologs often contain N-terminal domains found in polymorphic T6SS specialized effectors, such as PAAR (e.g., WP_047722894.1) or VgrG (e.g., WP_013660774.1). The gene downstream of *psi1* (A7P61_RS17170) encodes a member of the Tai4 family of immunity proteins (Russel *et al*., 2012). Interestingly, homologs of this gene are present immediately downstream of genes encoding Tae4 antibacterial effectors within the T6SS1 *hcp* island of other *Pantoea* strains (e.g., WP_182686054.1 in *Pantoea agglomerans* strain RSO7). The gene downstream of *tai4* (A7P61_RS17165) is a small open reading frame (ORF) similar to COG3895 domains (88.89% probability of similarity between A7P61_RS17165 amino acids 2-99 and amino acids 3-91 of the COG3895 domain consensus according to HHpred; Gabler *et al*., 2020), which are annotated as inhibitors of C-type lysozyme. Homologs of this gene are found in *Pantoea hcp* islands immediately downstream of a gene encoding a putative antibacterial effector containing endolysin, a peptidoglycan-targeting toxin domain (e.g., WP_115064794.1 in *Pantoea agglomerans* strain LMAE-2); therefore, this gene possibly encodes an immunity protein. The two genes upstream of *tagH* (A7P61_RS17160 and A7P61_RS17155) both encode proteins containing the lysozyme inhibitor LprI domain (according to NCBI Conserved Domain Database; Marchler-Bauer *et al*., 2007), homologs of which are found in *Pantoea hcp* islands immediately downstream of an antibacterial effector containing a lysozyme-like toxin domain (e.g., WP_090962028.1 in *Pantoea* sp. OV426), and could therefore encode immunity proteins. These observations indicate that the *Pab hcp* island contains a pore-forming toxin and its cognate immunity, as well as four orphan immunity proteins that were possibly maintained or acquired to provide immunity against peptidoglycan-targeting effectors deployed by competing *Pantoea* strains.

#### *vgrG* island

The *Pab vgrG* island (Fig. 1; Supplementary Fig. S1) includes a gene encoding a specialized VgrG effector (A7P61_RS17105) that harbors a C-terminal extension domain predicted to be a member of the glucosaminidase superfamily of peptidoglycan-targeting toxins. We predict that the gene downstream of the *vgrG* encodes its cognate immunity protein, which we termed Vgi (<u>V</u>grG <u>g</u>lucosaminidase immunity). The gene downstream of VgrG and its putative cognate immunity (A7P61_RS25185) is a homolog of genes found immediately downstream of specialized VgrG effectors in other *Pantoea* strains that contain a C-terminal glycosyl hydrolase 108 toxin domain (e.g., WP_111142272.1 in *Pantoea* sp. ARC607), and is therefore predicted to encode an immunity protein. We were unable to predict a role for the protein encoded by the A7P61_RS17090 gene.

#### *paar* island

The gene upstream of the gene encoding the PAAR domain-containing protein (hereafter, referred to as PAAR; A7P61_RS17085) encodes a protein with a DcrB domain (Fig. 1; Supporting Information Fig. S1). This domain was previously referred to as DUF1795 or Eag and it serves as an adaptor for T6SS specialized effectors (Alcoforado Diniz and Coulthurst, 2015; Cianfanelli *et al*., 2016). In addition to a PAAR domain, the PAAR-encoding gene (A7P61_RS17080) also encodes RhsA repeats and a C-terminal extension that harbors a putative antibacterial toxin domain. Homologs of this C-terminal domain are found in diverse *Proteobacteria*, where they are fused to PAAR and RhsA domains, and in *Firmicutes*, where they are fused to TANFOR domains, which were recently shown to carry polymorphic toxins (e.g., WP_066428025) (Quentin *et al*., 2018; Jana *et al*., 2020). Therefore, the PAAR protein appears to be a specialized effector. In agreement with the prediction that the PAAR C-terminal domain is a putative antibacterial toxin, a small gene (A7P61_RS25180), which potentially encodes a cognate immunity protein, is found immediately downstream to it.

Close inspection of the 3’ end of the T6SS1 gene cluster led us to conclude that the previous assessment of the *Pantoea* T6SS1 cluster borders (De Maayer *et al*., 2011) was incomplete; we determined that the cluster does not end immediately after the PAAR-encoding gene, but rather it extends downstream to the A7P61_RS17020 gene. We termed the region spanning from the adaptor-encoding gene upstream of the PAAR protein (A7P61_RS17085) to the A7P61_17020 gene as “*paar* island” (Fig. 1; Supporting Information Fig. S1). Analysis of the genes present in this island revealed several putative orphan toxin-immunity modules; this phenomenon was previously described downstream of other Rhs-containing polymorphic toxins (Koskiniemi *et al*., 2013). The A7P61_RS24260 gene encodes a small protein containing a TNT toxin domain; homologs of the TNT toxin are found fused to a PAAR homolog in other *Pantoea* strains (e.g., WP_201500397 in *Pantoea agglomerans* strain GR13). Downstream of the TNT toxin is a pseudogene (position 657,724-657,348 in the NZ**_**LXSW01000076.1 RefSeq assembly), which is not annotated in the *Pab* RefSeq genome assembly probably because of an apparent frame shift within the ORF. A homolog of this pseudogene is found immediately downstream of the above mentioned PAAR-fused TNT toxin in other *Pantoea* strains. Therefore, these two genes appear to encode an orphan toxin-immunity module. The A7P61_RS17070 gene encodes a homolog of proteins encoded immediately downstream of PAAR proteins at the same position in clusters of other *Pantoea* strains (e.g., WP_132498480.1 in *Pantoea agglomerans* strain MM2021_7), and therefore it is possibly an orphan immunity protein. The A7P61_RS24955 gene, which is annotated as a pseudogene, appears to encode a Rhs-containing protein that is missing its N-terminal region. The downstream A7P61_RS17055 gene encodes a homolog of proteins encoded immediately downstream of PAAR proteins at the same position in clusters of other *Pantoea* strains (e.g., WP_085068093.1 in *Pantoea alhagi* strain LTYR-11Z). Therefore, these two genes possibly represent another orphan toxin-immunity module that can be fused via DNA rearrangement to the PAAR encoded at the beginning of the *paar* island. The A7P61_RS17045 gene encodes another putative orphan immunity protein, as it is identical to a putative immunity protein encoded immediately downstream of a Rhs-containing PAAR protein that is at the same position of the *Pab* PAAR in the *paar* island, but in a different *Pantoea* strain (e.g., WP_010246002.1 in *Pantoea agglomerans* strain L15). Similar to A7P61_RS24955, the A7P61_RS24950 gene, which is also annotated as a Rhs-encoding pseudogene, appears to encode a protein missing both its N-terminal region and toxin-containing C-terminal region. Interestingly, the downstream A7P61_RS25175 gene encodes a protein displaying homology to the C-terminus of Rhs-containing PAAR proteins of other *Pantoea* strains (e.g., WP_062757477.1 in *Pantoea agglomerans* NBRC 102470), and is therefore possibly a toxin. As expected, the downstream A7P61_RS17035 gene encodes a homolog of the protein encoded downstream of the above-mentioned toxin that is predicted to be its cognate immunity protein. The A7P61_RS17030 gene may also encode an orphan immunity protein since it is homologous to a protein encoded immediately downstream of a Rhs-containing gene, which is found downstream of a Rhs-containing PAAR in other bacteria (e.g., WP_013036149.1 in *Erwinia amylovora* CFBP1430). The last two genes in this island (A7P61_RS17025 and A7P61_RS17020, which we termed *pse2* and *psi2*, respectively, as explained below), encode proteins that do not contain any described domain; however, their organization in a bicistronic operon and their position within the *paar* island suggest that they encode either an effector and immunity pair, or an orphan toxin-immunity module. The latter possibility is less likely since we did not identify homologs of the putative effector, which is encoded by the A7P61_RS17025 gene, fused to known N-terminal domains of polymorphic toxins.

Taken together, these observations indicate that the *Pab* T6SS1 cluster contains three rapidly evolving islands that accumulate arrays of orphan immunity and toxin-immunity modules. This possibly confers *Pab* the ability to resist aggression by *Pantoea* relatives that carry diverse T6SS effector repertoires, as well as allows *Pab* to evolve new effectors of its own by replacing the C-terminal toxin domain fused to the PAAR protein with toxin-immunity module present in the array downstream to it.

### Pab *T6SS1 is a functional antibacterial system*

To determine whether the T6SS1 cluster encodes a functional T6SS, we set out to monitor secretion of the hallmark secreted tail tube component Hcp (Mougous *et al*., 2006). To this end, we first constructed a T6SS mutant by deleting the gene encoding the conserved structural component TssA (Planamente *et al*., 2016; Zoued *et al*., 2016). As shown in Fig. 2A, exogenously expressed FLAG-tagged Hcp was secreted from wild-type *Pab* grown under standard laboratory conditions (i.e., LB at 28°C), but not from the Δ*tssA* mutant. Complementation of TssA from a plasmid restored Hcp secretion. This result demonstrates that the *Pab* T6SS1 is functional under laboratory conditions.

**Fig. 2.**
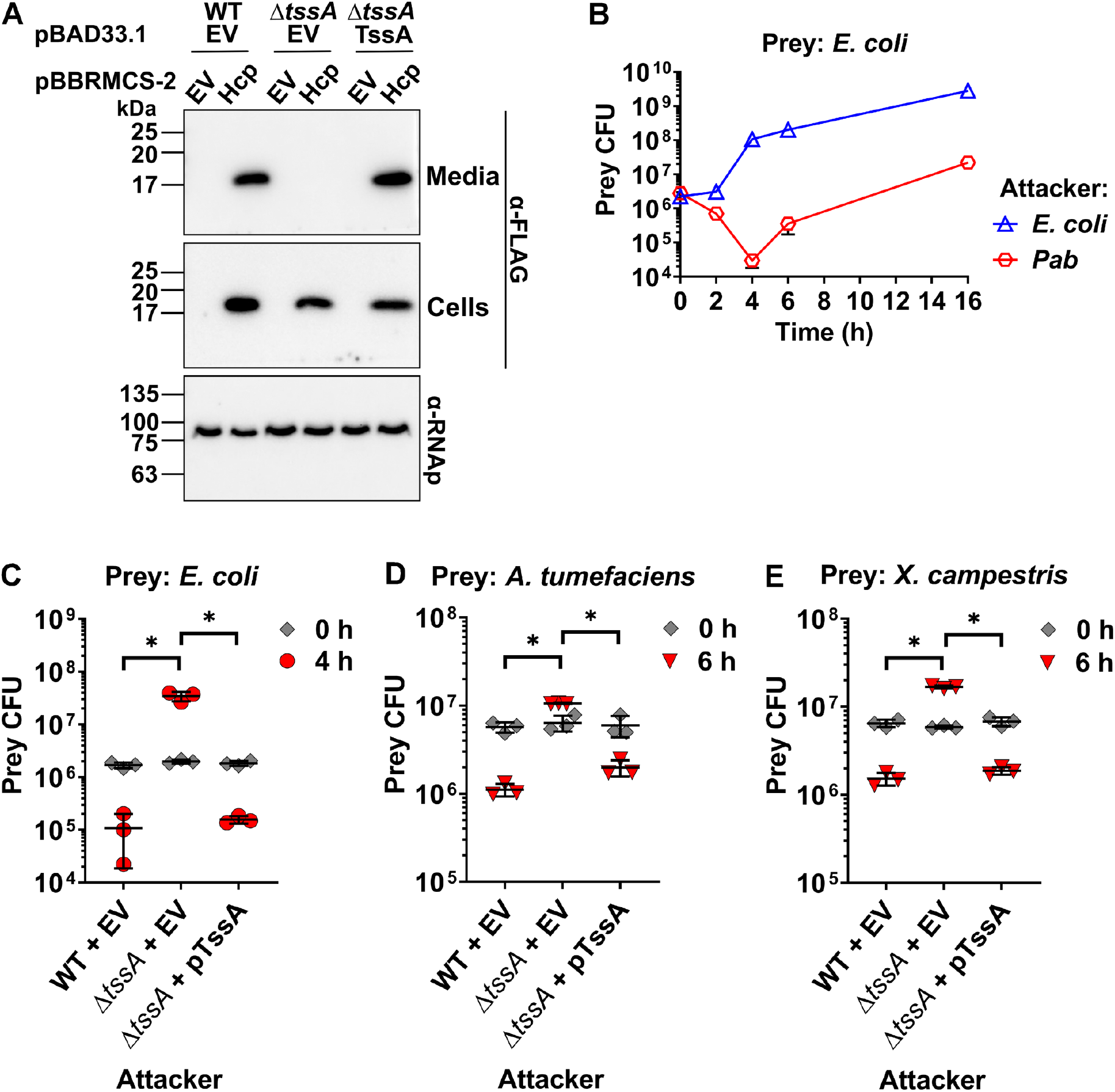
The *Pab* T6SS is a functional antibacterial system. **A**. Expression (cells) and secretion (media) of FLAG-tagged Hcp from *Pab* wild-type (WT) and Δ*tssA* mutant (Δ*tssA*) strains carrying the indicated plasmids were detected by immunoblotting using α-FLAG antibodies. RNA polymerase β was used to confirm equal loading in cells samples and was detected with α-RNAp antibodies. TssA was expressed from the arabinose-inducible plasmid, pBAD33.1. The experiment was repeated three times with similar results. Results from a representative experiment are shown. **B**. Viability counts of *E. coli* MG1655 prey at the indicated timepoints when co-incubated with either *E. coli* or *Pab* attackers. **C-E**. Viability counts of the preys *E. coli* MG1655 (C), *Agrobacterium tumefaciens* GV3101 (D), and *Xanthomonas campestris* pv. *campestris* 8100 (E) before (0 h) and after (4 h for *E. coli*; 6 h for *A. tumefaciens* and *X. campestris*) co-incubation with *Pab* WT and *ΔtssA* attackers carrying a plasmid either empty (EV) or expressing TssA. Data are shown as the mean ± SD of three biological replicates. Asterisks denote statistical significance between samples at t=4 h or t=6 h by an unpaired, two-tailed Student’s *t*-test (*P* < 0.01).

Our analysis of the T6SS1 gene cluster revealed several putative antibacterial effector and immunity pairs. Therefore, we hypothesized that this system plays a role in interbacterial competition. To test this hypothesis, we determined the outcome of co-incubation of *Pab* with *E. coli* MG1655 as potential sensitive prey. As shown in Fig. 2B, the viability of the *E. coli* prey decreased over time upon co-incubation with wild-type *Pab*, but not when *E. coli* was co-incubated with the equivalent numbers of *E. coli* attackers (*E. coli* prey contained a selective marker distinguishing them from their parental *E. coli* attackers); the decrease in *E. coli* viability reached a maximum after 4 h of co-incubation with *Pab*, followed by an increase in the *E. coli* prey CFU, which could be attributed to phase separation that protected pockets of *E. coli* populations (McNally *et al*., 2017). To determine whether the loss of *E. coli* viability was mediated by the *Pab* T6SS1, we co-incubated *E. coli* prey with either wild-type *Pab* or the Δ*tssA* mutant, and monitored its viability after 4 h. As expected, loss of *E. coli* viability was detected when it was co-incubated with wild-type *Pab* but not with the Δ*tssA* mutant (Fig. 2C), indicating that T6SS1 is responsible for the antibacterial effect. Deletion of the *tssA* gene did not affect *Pab* bacterial growth (Supporting Information Fig. S2). Notably, exogenous complementation of TssA from a plasmid restored the antibacterial activity of the Δ*tssA* mutant (Fig. 2C).

Since *E. coli* is not commonly found in the *Pab* natural environmental niche, the ryzosphere (i.e., soil and plant roots), we tested whether the *Pab* T6SS1 can also mediate the killing of plant-associated bacteria. To this end, we monitored the viability of *Agrobacterium tumefaciens* and *Xanthomonas campestris* pv. *campestris* upon co-incubation with *Pab*. As shown in Fig. 2D and E, *Pab* efficiently killed both prey bacteria. This antibacterial activity was mediated by T6SS1, since it was abolished when the Δ*tssA* mutant was used (Fig. 2D and E). Taken together, these results demonstrate that the *Pab* T6SS1 is an antibacterial system that is used to eliminate bacterial competitors.

### VgrG is a specialized antibacterial effector targeting the peptidoglycan

We set out to investigate antibacterial effector and immunity pairs that are encoded within the T6SS1 gene cluster. We first focused on VgrG and its cognate immunity Vgi. As mentioned above, the *Pab* VgrG is a specialized effector that contains a C-terminal extension domain predicted to be a member of the glucosaminidase superfamily (Fig. 3A; amino acids 675-829). The presence of this domain suggests that the VgrG effector hydrolyses the peptidoglycan in the bacterial periplasm. In agreement with this prediction, the putative cognate immunity protein Vgi, which is encoded by the gene downstream of *vgrG*, has an N-terminal signal peptide for periplasmic localization (according to the SignalP 5.0 prediction server; Almagro Armenteros *et al*., 2019) (Fig. 3A). To investigate whether VgrG is an antibacterial toxin, we expressed it in *E. coli* as a surrogate host from an arabinose-inducible plasmid, either in its natural form (_cyto_VgrG) or fused to an N-terminal PelB signal peptide (_peri_VgrG), for periplasmic localization. As shown in Fig. 3B, only the periplasmic version was toxic in agreement with its predicted activity as a peptidoglycan hydrolyzing toxin. Induction of _peri_VgrG expression resulted in a decline in the OD_600_ readings, which is compatible with cell lysis.

**Fig. 3.**
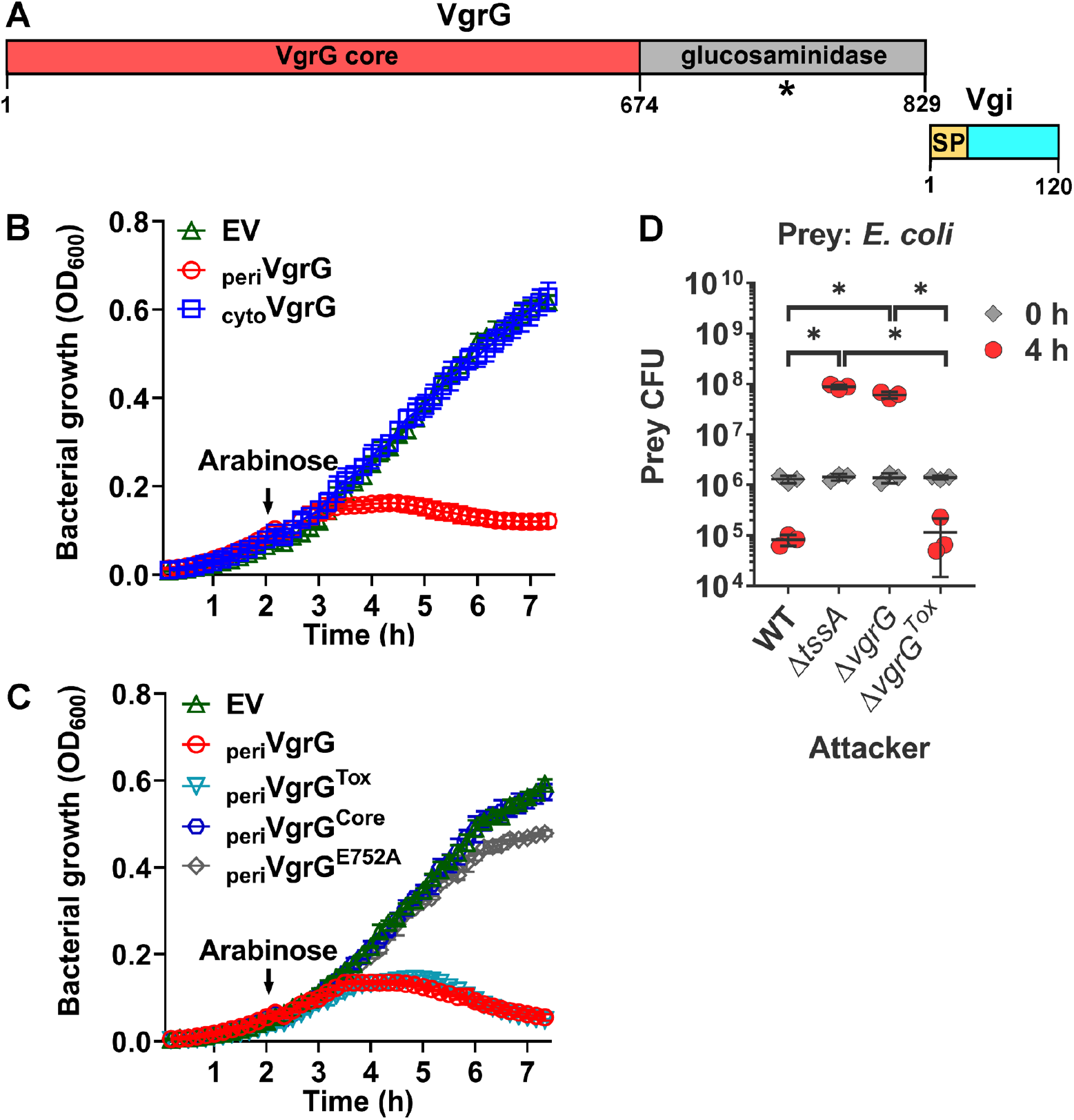
The C-terminus of VgrG contains an antibacterial toxin. **A**. Schematic representation of *Pab* VgrG and its cognate immunity protein Vgi. Domains and amino acid positions of note are denoted. An asterisk indicates the position of the conserved E752 amino acid residue. SP, signal peptide. **B-C**. Growth of *E. coli* MG1655 containing arabinose-inducible plasmids for the expression of cytoplasmic (cyto) or periplasmic (peri) VgrG (B), and periplasmic VgrG derivatives (C), including the VgrG glucosaminidase domain (amino acids 675-829; _peri_VgrG^Tox^), the VgrG core (amino acids 1-674; _peri_VgrG^Core^), and VgrG carrying a substitution of the conserved E752 to Ala (_peri_VgrG^E752A^). Protein expression was induced after 2 h of growth by the addition of 0.05% L-arabinose (denoted by an arrow). Data are shown as the mean ± SD (n=4 technical replicates) from a representative experiment. Experiments were repeated three times with similar results. **D**. Viability counts of *E. coli* MG1655 prey before (0 h) and after (4 h) co-incubation with the attackers *Pab* wild-type (WT), *Pab ΔtssA* mutant (*ΔtssA*), *Pab* Δ*vgrG* mutant (Δ*vgrG*), or *Pab* with a deletion in the region encoding the VgrG glucosaminidase domain (Δ*vgrG*^*Tox*^). Data are shown as the mean of three biological replicates. Asterisks denote statistical significance between samples at t=4 h by an unpaired, two-tailed Student’s *t*-test (*P* < 0.01).

Next, we determined whether the C-terminal glucosaminidase domain is responsible for the toxicity observed when VgrG was delivered to the *E. coli* periplasm. Indeed, expression of the glucosaminidase domain (amino acids 675-829; _peri_VgrG^Tox^), but not of the N-terminal structural part of VgrG (amino acids 1-674; _peri_VgrG^Core^), was sufficient to cause the observed decline in the OD_600_ of the *E. coli* culture (Fig. 3C). Analysis of protein sequences homologous to the VgrG C-terminal toxin domain (Supporting Information Fig. S3) revealed a conserved glutamic acid (E752) that corresponds to an annotated active site residue in another member of the glucosaminidase family, the peptidoglycan-hydrolyzing flagellar component, FlgJ (UniProtKB – P75942; Boutet *et al*., 2007). In agreement with this C-terminal domain being the toxin, and with its predicted activity as a member of the glucosaminidase superfamily, substitution of the conserved E752 to alanine (_peri_VgrG^E752A^) abolished the toxicity of VgrG in *E. coli* (Fig. 3C).

The expression of all VgrG versions in *E. coli* was confirmed by immunoblotting, aside from the toxic C-terminal domain alone (Supporting Information Fig. S4). To further confirm that the C-terminal glucosaminidase domain of VgrG is a toxin domain that is not required for the structural role of VgrG as an essential part of the T6SS tail tube, we constructed *Pab* mutant strains in which we deleted either the entire *vgrG* gene (Δ*vgrG*), or only the region encoding the C-terminal glucosaminidase domain (Δ*vgrG*^*Tox*^). We then used these mutants as attackers in a competition assay against *E. coli* prey. Deletion of the entire *vgrG* gene abolished T6SS-mediated antibacterial activity, similar to deletion of the conserved component *tssA* (Fig. 3D). However, deletion of the C-terminal toxin domain alone did not affect the antibacterial activity against *E. coli* prey, indicating that the T6SS remained functional and delivered other antibacterial effectors even in the absence of the specialized toxin domain fused to VgrG.

The toxicity of VgrG in the *E. coli* periplasm and the reduction in the optical density of the *E. coli* culture upon its expression (Fig. 3B) supported the predicted activity of the C-terminal toxin domain as a peptidoglycan hydrolase. To demonstrate this predicted activity, we monitored the effect of periplasmic expression of VgrG on cell morphology and the peptidoglycan layer of *E. coli* under the microscope. Expression of VgrG induced cell rounding and lysis, together with apparent disappearance of the peptidoglycan layer (Fig. 4 top panel; Supporting Information movie S1). The catalytically inactive VgrG mutant (VgrG^E752A^) had no effect on cell morphology or the peptidoglycan over time (Fig. 4 middle panel; Supporting Information movie S2), and it was comparable to an empty expression plasmid (Fig. 4 bottom panel; Supporting Information movie S3). Taken together, these results demonstrate that VgrG is a specialized antibacterial effector with a C-terminal glucosaminidase toxin domain that induces peptidoglycan hydrolysis leading to cell lysis. Interestingly, we found that homologs of VgrG found in other *Pantoea* strains contain one of at least five C-terminal toxin domains. Aside from the glucosaminidase domain present in the *Pab* VgrG, other homologs contain LytD, glycosyl hydrolase 108, lysozyme-like, or metallopeptidase C-terminal domains (Supporting Information Fig. S5). Remarkably, these five domains are all predicted to target the peptidoglycan.

**Fig. 4.**
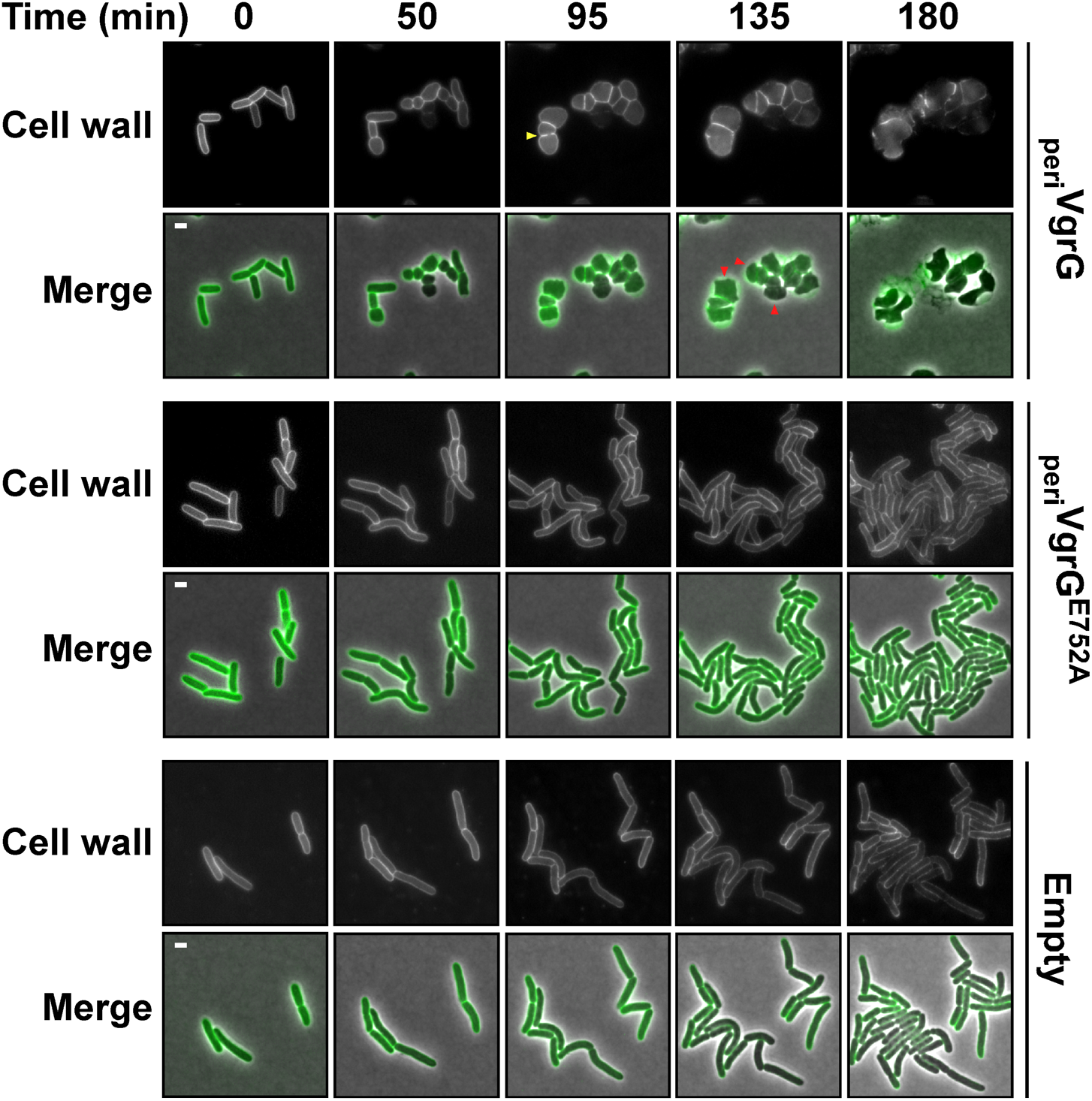
VgrG induces cell wall degradation leading to cell lysis in *E. coli*. Time-lapse microscopy of *E. coli* cells expressing periplasm-targeted VgrG (_peri_VgrG), a catalytic mutant (_peri_VgrG^E752A^), or containing an empty arabinose-inducible expression vector. Cells stained with Wheat Germ Agglutinin Alexa Fluor 488 Conjugate (cell wall stain; green in merged panels) were spotted on LB agarose pads supplemented with kanamycin and 0.5% arabinose. Merging of the phase contrast and GFP channels (for Alexa Fluor 488 visualization) and only the GFP channel are shown. A yellow arrowhead denotes cell wall stain that disappears in the following time frame. Red arrowheads denote cells that are lysed in the following time frame. Scale bar=2 μm.

### Vgi antagonizes the toxicity mediated by VgrG

We predicted that the gene downstream of *vgrG*, encoding Vgi, is its cognate immunity gene. As expected from an immunity against a toxin that functions in the periplasm, Vgi has a predicted signal peptide. To investigate the ability of Vgi to antagonize VgrG-mediated toxicity, we first expressed both proteins, either together or alone, in *E. coli*. As shown in Fig. 5A, Vgi rescued *E. coli* from VgrG-mediated toxicity when periplasm localized VgrG and Vgi were co-expressed. In addition, we tested whether VgrG and Vgi function as a T6SS antibacterial effector and immunity pair. To this end, we generated a *Pab* mutant in which we deleted the *vgrG* region encoding the C-terminal toxin domain together with the *vgi* gene (Δ*vgrG*^*Tox*^*-vgi*). This deletion did not affect *Pab* growth (Supporting Information Fig. S2). We then used this mutant strain as prey in competition against *Pab* attackers. The deletion rendered the mutant susceptible to intoxication by a wild-type *Pab* attacker, but not by the T6SS1 deficient attackers Δ*tssA* and Δ*vgrG* (Fig. 5B), indicating that a gene providing immunity against T6SS-mediated attack was deleted in this prey. A Δ*vgrG*^*Tox*^ attacker, which still retains ability to intoxicate *E. coli* prey (Fig. 3D), was unable to intoxicate the mutant *Pab* prey, demonstrating that the intoxicating effector was the C-terminal toxin domain in the specialized VgrG. Importantly, exogenous expression of Vgi from an arabinose-inducible plasmid restored the ability of the Δ*vgrG*^*Tox*^*-vgi* prey to resist intoxication by a wild-type *Pab* attacker (Fig. 5B). Therefore, the results show that VgrG and Vgi are a *bona fide* T6SS antibacterial effector and immunity pair.

**Fig. 5.**
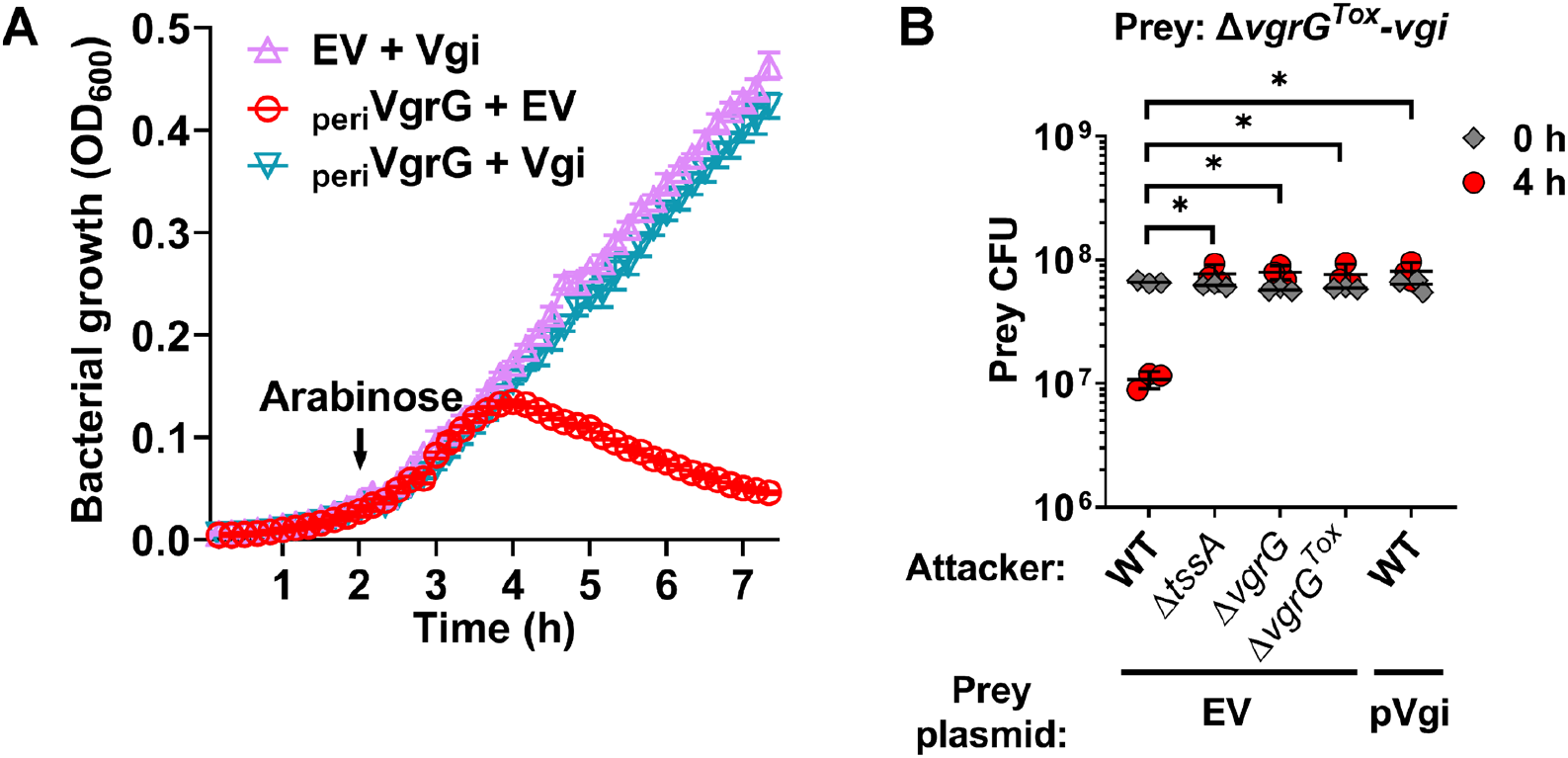
Vgi antagonizes VgrG-mediated toxicity. **A**. Growth of *E. coli* MG1655 containing arabinose-inducible plasmids for co-expression of periplasmic VgrG (_peri_VgrG) and Vgi. Empty vectors (EV) were used as control. Protein expression was induced after 2 h of growth by the addition of 0.05% L-arabinose (denoted by an arrow). Data are shown as the mean±SD, n=4 technical repeats. **B**. Viability counts of *Pab* mutant lacking the VgrG glucosaminidase domain and the downstream *vgi* gene (*ΔvgrG*^*Tox*^*-vgi*), and carrying a plasmid either empty (EV) or for arabinose-inducible expression of Vgi, before (0 h) and after (4 h) co-incubation with the indicated *Pab* attackers on LB agar containing 0.05% L-arabinose. Data are shown as the mean of three biological replicates. Asterisks denote statistical significance between samples at t=4 h by an unpaired, two-tailed Student’s *t*-test (*P* < 0.01).

### Pse2 is a T6SS-delivered antibacterial effector that is antagonized by Psi2

In the analysis of the *Pab* T6SS1 gene cluster, we hypothesized that the two genes at the 3’ end of the cluster, encoding Pse2 and Psi2, are a T6SS effector and immunity pair. To test this hypothesis, we first determined whether Pse2 is secreted in a T6SS-dependent manner. Immunoblotting of ectopically expressed Pse2 revealed its presence in the supernatant of wild-type *Pab*, but not in the supernatant of Δ*tssA Pab* (Fig. 6A). This result confirms that Pse2 is a secreted substrate of T6SS1.

**Fig. 6.**
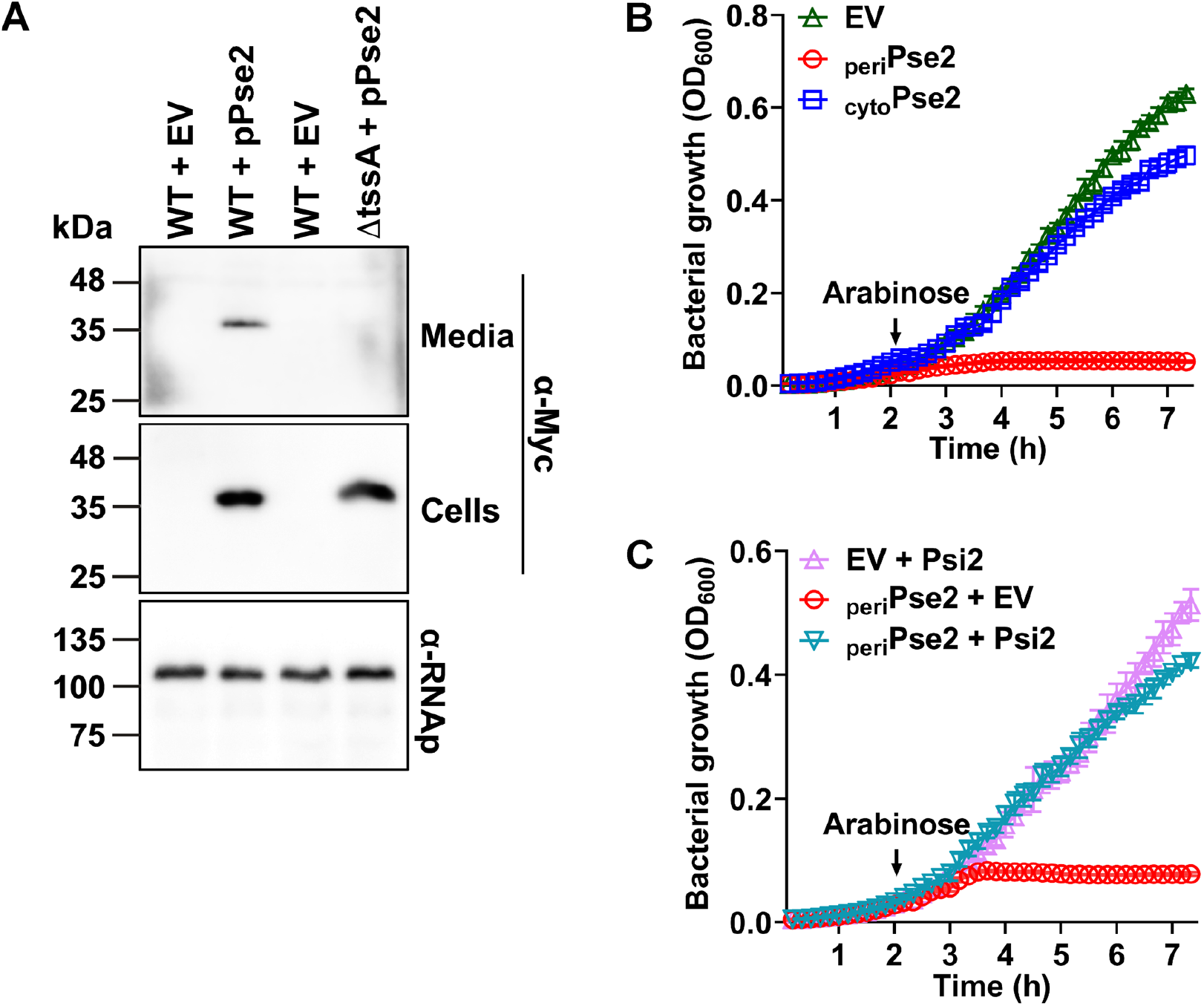
Pse2 and Psi2 are an antibacterial effector and immunity pair. **A**. Expression (cells) and secretion (media) of Pse2 from *Pab* wild-type (WT) and Δ*tssA* mutant (Δ*tssA*) strains carrying a plasmid for arabinose-inducible expression of C-terminal Myc-tagged Pse2 or an empty vector (EV). Pse2-Myc was detected by immunoblotting using α-Myc antibodies. RNA polymerase β was used to confirm equal loading in cell samples and was detected with α-RNAp antibodies. **B-C**. Growth of *E. coli* MG1655 containing arabinose-inducible plasmids for the expression of cytoplasmic (cyto) or periplasmic (peri) Pse2 (B), or periplasmic Pse2 together with Psi2 (C). Empty vectors (EV) were used as control. Protein expression was induced after 2 h growth by the addition of 0.05% L-arabinose (denoted by an arrow). Data are shown as the mean±SD, n=4 technical repeats. In A-C, experiments were repeated three times with similar results. Results from a representative experiment are shown.

Both Pse2 and Psi2 contain predicted transmembrane helices (according to Phobius webserver predictions; Käll *et al*., 2004) (Supporting Information Fig. S6A), suggesting that the putative effector Pse2 functions in the bacterial membrane or periplasm. To determine whether Pse2 mediates antibacterial toxicity, we exogenously expressed it from an arabinose-inducible plasmid in *E. coli*, either in its natural form (_cyto_Pse2) or fused to an N-terminal PelB signal peptide (_peri_Pse2), for periplasmic localization. As shown in Fig. 6B, only the periplasmic version was toxic. Notably, expression of Pse2 in the periplasm did not result in a decline in OD_600_ readings over time; rather, OD_600_ readings remained stable, suggesting that the toxic effect of Pse2 is bacteriostatic. The expression of the cytoplasmic version of Pse2, but not of its periplasmic version, was confirmed by immunoblotting (Supporting Information Fig. S6B). It is possible that due to its toxicity, the accumulation of the periplasmic Pse2 was below detection limit. Furthermore, co-expression of Psi2 rescued *E. coli* from Pse2-mediated toxicity (Fig. 6C). Taken together, these results support the hypothesis that Pse2 and Psi2 are a *Pab* T6SS1 antibacterial effector and immunity pair.

## DISCUSSION

We identified a functional T6SS cluster (T6SS1) in the genome of *Pab* phytopathogenic bacteria mediating antibacterial activity and enabling *Pab* to outcompete several bacterial species that co-inhabit the rhizosphere niche. *Pab* also contains an auxiliary T6SS gene cluster, named T6SS2, encoding additional copies of T6SS core and accessory proteins. Clusters with a genetic composition similar to *Pab* T6SS1 and T6SS2 have been previously reported to be conserved in the genome of *Pantoea ananatis* and *Erwinia* species, while an additional cluster, which is named T6SS3 and present in a subset of *Pantoea* and *Erwinia* strains, is absent in *Pab* (De Maayer *et al*., 2011; Shyntum *et al*., 2014). The role of T6SS components found in the partial T6SS2 cluster, and whether or not the corresponding genes are expressed, remain to be investigated. Since the gene content and organization of the T6SS2 cluster appear to be conserved in *Pab* and *P. ananatis* strains (Shyntum *et al*., 2014), it is likely that T6SS2 serves a functional purpose favoring its retention in the genus. It is possible that T6SS2 and T6SS1 components function together to form a derivative T6SS apparatus.

We identified antibacterial effector and immunity gene pairs encoded in three dynamic islands (*hcp, vgrG* and *paar*) within the T6SS1 gene cluster. A distinctive characteristic of the composition of the *hcp* island is the presence of an effector immunity gene pair downstream of the *hcp* gene followed by an array of putative orphan immunities. This gene organization possibly evolved as a result of consecutive integration events of effector and cognate immunity genes by a yet to be defined mechanism of horizontal gene transfer between *Pantoea* and *Erwinia* strains. We hypothesize that integration of each new module replaced a previous effector gene, while the previous immunity gene was retained downstream of the newly integrated cassette. Therefore, the array of orphan immunity genes present in the *hcp* island may be originated by multiple events of effector and immunity integration and effector displacement, possibly maintaining immunity against attacks by other kin strains in which the replaced effectors are still functional. Similar genetic rearrangements were previously proposed as the evolutionary dynamics that generated arrays of multiple different orphan immunity genes in T6SS clusters of *V. cholerae* strains (Kirchberger *et al*., 2017). Similar dynamics can be also envisaged for the origin and function of the orphan immunity gene present in the *vgrG* island downstream of the *vgrG* and cognate immunity genes.

An alternative mechanism of effector diversification can be found in the *paar* island. The conserved effector in this island is a PAAR domain-containing protein that harbors Rhs repeats and a toxin domain fused to the C-terminal end of the protein; the toxin domain differs between *Pantoea* strains. Similar to previously reported Rhs proteins (Ma *et al*., 2016), the downstream regions of the *Pantoea* Rhs effectors harbor various orphan toxin and immunity modules that have the potential to recombine into the main PAAR-Rhs-encoding gene, thus giving rise to different specialized effectors than can outcompete their parental kin. Notably, some of the orphan toxin and immunity modules in *Pab paar* islands appear to have been deactivated by frame-shift or nonsense mutations (i.e., pseudogenes). Taken together, these observations indicate that the *Pab* T6SS1 cluster is rapidly evolving and likely to drive strain diversification. We propose that in a sense, the *hcp* and *vgrG* islands record the evolutionary effector history of the strain, whereas the *paar* island contains information of possible future effectors.

The *vgrG* gene of *Pab* bacteria encodes an evolved form of the VgrG tail component with a C-terminal glucosaminidase extension. This extension is missing in *P. ananatis* strains with a single *vgrG* copy, but present in an additional *vgrG* gene of *P. ananatis* strains with two *vgrG* copies (Shyntum *et al*., 2014). Based on computational analyses, we predicted that the VgrG C-terminal domain encodes a peptidoglycan-hydrolyzing toxin that targets a periplasmic component of prey cells. Indeed, a combination of genetic, toxicity, and microscopy assays confirmed this hypothesis. Notably, the presence of peptidoglycan-hydrolyzing toxin domains in homologs of VgrG identified in other *Pantoea* strains suggests that the role of this effector in targeting the cell wall of a competing cell is an important aspect of this T6SS, and this activity is thus conserved.

We found a novel effector and immunity pair, Pse2/Psi2, which is encoded at the 3’ end of the *paar* island. Although the activity and specific target(s) of the antibacterial effector Pse2 remain unknown, the presence of transmembrane helices in its amino acid sequence suggests that it could target the bacterial membrane, possibly by forming pores. It is also probable that other effectors are encoded outside of the *Pab* T6SS1 cluster, and that these can further diversify its antibacterial arsenal.

In conclusion, we identified a functional antibacterial T6SS and several effector and immunity pairs in *Pab* phytopathogenic bacteria. Future studies will be aimed to assess the contribution of *Pab* antibacterial activity to competitive fitness in the rhizosphere, and whether this activity contributes directly or indirectly to *Pab* virulence towards the host plant.

## EXPERIMENTAL PROCEDURES

### Bacterial strains and media

The bacterial strains used are: *Pantoea agglomerans* pv. *betae* strain 4188 (*Pab*), *Agrobacterium tumefaciens* GV3101 (Han *et al*., 2013), *Xanthomonas campestris* pv. *campestris* strain B100 (Hötte *et al*., 1990), *Escherichia coli* strains DH5α *λpir* (Miller and Mekalanos 1988; Penfold and Pemberton 1992) and MG1655 (Edwards and Palsson, 2000). *P. agglomerans, A. tumefaciens*, and *X. campestris* were grown in Lysogeny Broth (LB) medium at 28°C; *E. coli* were grown in LB or 2xYT medium at 37°C. Media were supplemented with ampicillin (50 μg/mL), spectinomycin (50 μg/mL), rifampicin (100 μg/mL), streptomycin (50 μg/mL), tetracycline (13 μg/mL) or chloramphenicol (35 μg/mL), as required.

### T6SS cluster analysis

*Pab* T6SS gene clusters were identified by homology searches using T6SS components that were previously reported in other *Pantoea* strains (De Maayer *et al*., 2011). Analyses of domains encoded by genes located in the *hcp, vgrG* and *paar* islands were performed using the NCBI Conserved Domain Database (Lu *et al*., 2020) and hidden Markov modelling (HHpred; Zimmermann *et al*., 2018). Homologs in other bacterial strains were identified using BLAST (Altschul *et al*., 1990), and their genomic neighborhoods were analyzed manually.

### VgrG^Tox^ amino acid conservation

One round of PSI-BLAST (Altschul *et al*., 1997) was used to identify 500 proteins homologous to the VgrG C-terminal toxin domain (amino acids 675-829; VgrG^Tox^). The retrieved sequences were aligned in MEGA X (Kumar *et al*., 2018) using MUSCLE (Edgar, 2004). Aligned columns not found in VgrG were discarded. The amino acid conservation of the homologs was illustrated using the WebLogo 3 server (http://weblogo.threeplusone.com; Crooks *et al*., 2004). The putative active site of the VgrG toxic domain was determined as E752 based on amino acid sequence comparison with the *Salmonella typhimurium* FlgJ peptidoglycan hydrolase (Zaloba *et al*., 2016).

### Pse2 amino acid conservation

Three rounds of PSI-BLAST (Altschul *et al*., 1997) were used to identify 500 proteins homologous to Pse2. The retrieved sequences were aligned in MEGA X (Kumar *et al*., 2018) using MUSCLE (Edgar, 2004). Aligned columns not found in Pse2 were discarded. The amino acid conservation of the homologs was illustrated using the WebLogo 3 server (http://weblogo.threeplusone.com; Crooks *et al*., 2004). Secondary structures were predicted with Jpred4 (Drozdetskiy *et al*., 2015) using the Pse2 sequence; transmembrane helices were predicted with the Phobius server (Käll *et al*., 2004).

### Pab growth assays

*Pab* and its mutant derivative strains were grown overnight in LB at 28°C, diluted to OD_600_=0.01, and then 200 μL of each sample were transferred into 96-well plates in triplicates. OD_600_ values were measured every 10 min for 7 h while plates were grown at 28°C with agitation (205 cpm) in a microplate reader (BioTek SYNERGY H1).

### Plasmid construction

For expression in *E. coli*, genes encoding *tssA* (A7P61_RS17205), *vgrG* (A7P61_RS17105), *vgrG*^*Core*^, *vgrG*^*Tox*^, *vgi* (A7P61_RS17100), *pse2* (A7P61_RS17025), *psi2* (A7P61_RS17020), were amplified from *Pab* genomic DNA. By using the Gibson assembly method (Gibson *et al*., 2009), amplicons were inserted in the multiple cloning site (MCS) of one of the following arabinose-inducible plasmids: pBAD/*Myc*–6His (hereafter referred to as pBAD), or pPER5/*Myc*–6His (hereafter referred to as pPER5), a pBAD derivative carrying the PelB signal peptide sequence at the 5′ of the MCS (Dar *et al*., 2021), or pBAD33.1 (Chung and Raetz, 2010). All inserted genes were positioned in frame with a 3’-encoded *Myc*–6His tag. The obtained plasmids were transformed into *E. coli* MG1655 using the electroporation method.

For expression in *Pab*, the *hcp* (A7P61_RS17185), *tssA* (A7P61_RS17205), *pse2* (A7P61_RS17025) and *psi2* (A7P61_RS17020) genes were PCR-amplified from *Pab* genomic DNA and inserted in the MCS of the pBBR-MCS2 (Kovach *et al*., 1995) or pBAD plasmids. The obtained plasmids were transformed into *E. coli* DH5α *λpir* using the heat shock method (Froger and Hall, 2007), and then introduced into *Pab* bacteria by triparental mating using the conjugation helper strain *E. coli* DH5α carrying the helper plasmid pBR2073 (Figurski and Helinski, 1979). When an arabinose-inducible vector was used, transformants were grown on LB agar plates or liquid cultures supplemented by 0.4% (wt/vol) glucose to repress expression from the *Pbad* promoter.

### Site-directed mutagenesis

To generate the *vgrG*^*E752A*^ mutant, site-directed mutagenesis was performed using as template the *vgrG* gene cloned in the pPER5 plasmid, as described by Edelheit *et al*. (2009). *vgrG* was amplified in two separate reactions of asymmetric PCR using either a forward or reverse primer carrying a Glu to Ala substitution at position 752. Reaction products were combined, denaturated at 95°C, reannealed and digested by *Dpn*I restriction enzyme to remove methylated parental DNA. The product was then directly transformed in *E. coli* MG1655.

### Protein expression in E. coli

*E. coli* MG1655 cells carrying arabinose-inducible expression plasmids pBAD or pPER5 were grown overnight at 28°C in 3 mL of LB supplemented with appropriate antibiotics and 0.4% (wt/vol) glucose. Cultures were normalized to OD_600_=0.1 and grown for 2 h at 28°C. Cells were then washed with LB to remove residual glucose, and 0.1% (wt/vol) L-arabinose was added to the media to induce protein expression. Cultures were grown for 2 or 4 h at 28°C as necessary, and 0.25 OD_600_ units of cells were pelleted and resuspended in 40 μL (2X) Tris-Glycine SDS sample buffer (Novex, Life Sciences) supplemented with 0.05% (wt/vol) β-mercaptoethanol. Samples were boiled and cell lysates were fractionated by SDS-PAGE, transferred onto PDVF membranes, and immunoblotted with α-Myc (Santa Cruz Biotechnology, 9E10, mouse mAb) antibodies, used at 1:1,000 dilution.

### Bacterial toxicity assays

*E. coli* MG1655 cells transformed with arabinose-inducible expression plasmids pBAD or pPER5 harboring a kanamycin-resistance cassette (Fridman *et al*., 2020), or co-transformed with pPER5 and pBAD33.1 harboring a chloramphenicol-resistance cassette were grown overnight at 28°C in 2xYT supplemented with appropriate antibiotics and 0.4% (wt/vol) glucose to repress expression from the *Pbad* promoter. Cultures were washed with 2xYT to remove residual glucose and normalized to OD_600_=0.01 in 2xYT supplemented with the appropriate antibiotics. Next, 200 μL of each sample was transferred into 96-well plates in quadruplicates and grown at 28° in a BioTeck SYNERGY H1 microplate reader with agitation (205 cpm). After 2 h, L-arabinose was added to each well at a final concentration of 0.05% (wt/vol) to induce protein expression, and plates were re-inserted into the microplate reader for 5 h additional incubation. OD_600_ values were measured every 10 min.

### Generation of Pab deletion strains

To generate deletions of the *Pab tssA* and *vgrG* genes, and of the *vgrG* region encoding the C-terminal toxin domain either alone or together with its cognate immunity gene *vgi*, sequences located 1 kb upstream and 1 kb downstream of the region to be deleted, were PCR-amplified and inserted into the MCS of pDM4, a Cm^R^OriR6K suicide plasmid (Milton *et al*., 1996). The obtained plasmids were transformed into *E. coli* DH5α *λpir* by electroporation (Lessard, 2013). Transformants were used to transfer the construct into *Pab* by triparental mating using the helper strain *E. coli* DH5α carrying the pBR2073. A first selection of colonies was carried out on LB agar plates supplemented with chloramphenicol. Growing colonies were then transferred to LB agar plates supplemented with sucrose (15% [wt/vol]) for counter-selection and loss of the SacB-containing pDM4, as described (Salomon *et al*., 2013). Deletions were confirmed by PCR and sequencing.

### Bacterial competition assays

Attacker and prey bacterial strains were grown overnight in LB or 2xYT supplemented with the appropriate antibiotics. Bacterial cultures were normalized to OD_600_=0.5 (*Pab* and derivative strains) or OD_600_=0.4 (*E. coli, A. tumefaciens* and *X. campestris*) and mixed at a 4:1 (attacker:prey) volume ratio. Mixtures were spotted on LB agar plates with or without the addition of 0.1% (wt/vol) L-arabinose to induce expression from plasmids, and incubated at 28°. Colony-forming units (CFU) of prey spotted at t=0 h were determined by plating 10-fold serial dilutions on selective agar plates. To determine prey CFU at later timepoints, bacterial spots were scraped from agar plates into 1 mL LB. Next, 10-fold serial dilutions were plated on selective agar plates and CFU were calculated.

### Protein secretion assays

Protein secretion assays were performed as previously described (Fridman *et al*., 2020), with minor modifications. Briefly, *Pab* strains were grown overnight in LB supplemented with antibiotics, when required. The cultures were normalized to OD_600_=0.1 in 5 mL of LB supplemented with appropriate antibiotics and 0.005% (wt/vol) L-arabinose, and grown for 5 h at 28°C. For extraction of cellular proteins (cells fraction), 0.25 OD_600_ units were collected, and cell pellets were resuspended in 40 μL of reducing protein sample buffer (2X; 100 mM Tris/HCl pH= 7.0, 20% glycerol, 25% sodium dodecyl sulfate [SDS], bromophenol blue, 0.05% β-mercaptoethanol). The remaining culture volume (media fraction) was filtered (0.22 μm) and precipitated by addition of deoxycholate (150 μg/mL) followed by 8% trichloroacetic acid (Bensadoun and Weinstein, 1976). Precipitated proteins were washed twice with cold acetone and resuspended in 20 μL of 10 mM Tris/HCl pH=8.0. An equal volume of 2X protein sample buffer and 1 μL NaOH 5 N were added to each sample. Samples were boiled and then resolved on SDS-PAGE, transferred onto PDVF membranes and immunoblotted with α-Myc or α-FLAG (Santa Cruz Biotechnology, F1804, mouse mAb) antibodies used at a 1:1,000 dilution. RNA polymerase β (RNAp) was detected with Direct-Blot™ HRP α-*E. coli* RNA Sigma 70 (mouse mAb #663205; BioLegend; referred to as α-RNAp) and was used as a non-secreted protein to confirm equal loading (Dar *et al*., 2021).

### Microscopy

To determine the effect of protein expression in *E. coli*, overnight *E. coli* MG1655 cells carrying pPER5-based plasmids were diluted 100-fold into 3 mL of fresh LB supplemented with kanamycin 30 mg/mL and 0.4% (wt/vol) glucose. After 2 h incubation at 30°C, cells were washed twice with 0.15 M NaCl and normalized to OD_600_=1.5. For peptidoglycan visualization, cells were stained with Wheat Germ Agglutinin (WGA) Alexa Fluor 488 Conjugate (Biotium; Catalog no. 29022-1) at a final concentration of 0.1 mg/mL, and incubated for 10 min at room temperature (RT). Then, 1 μL of each culture was spotted onto LB agarose pads (1% [wt/vol] agarose supplemented with 30 mg/mL kanamycin and 0.2% [wt/vol] L-arabinose) to which 1 μL of the membrane-impermeable DNA dye, propidium iodide (PI;1 mg/mL; Sigma-Aldrich), was pre-applied. After the spots dried (1-2 minutes at RT), the agarose pads were mounted, facing down, onto 35 mm glass bottom CELLview™ cell culture dishes (Greiner). Cells were then imaged every 5 min for 3 h under a fluorescence microscope, as detailed below. The stage chamber (Okolab) temperature was set to 30°C. The following setup was used for imaging: a Nikon Eclipse Ti2E inverted motorized microscope with a CFI PLAN apochromat DM 100X oil lambda PH-3 (NA, 1.45) objective lens, a Lumencor SOLA SE II 395 light source, an ET-dsRED (#49005, CHROMA, to visualize PI signal) and ET-EGFP (#49002, CHROMA, to visualize GFP signal) filter sets, and a DS-QI2 Mono cooled digital microscope camera (16 MP). Obtained images were further processed and analyzed using Fiji ImageJ suite (Schindelin *et al*., 2019).

## ACNOWLEDGEMENTS

This work was supported by grants from the Israel Science Foundation (ISF; grant no. 488/19 to G.S. and I.B.; grant no. 920/17 to D.S.). C.M.F. was supported by scholarships from the Clore Israel foundation and from the Manna Center Program in Food Safety and Security at Tel Aviv University.

## SUPPORTING INFORMATION

**Fig. S1**. Evolutionary dynamics in *Pab* T6SS1 islands. Genes are represented by arrows indicating the direction of transcription. Gene names or locus tags (A7P61_RSxxxxx) in the *Pab* T6SS1 cluster are denoted above. Colors denote homology between genes found in different genomes. Associated effector or toxin (full arrows) and immunity (empty arrows) pairs are denoted by the same color.

**Fig. S2**. *Pab* and derivative strains display a similar growth rate. Growth of *Pab* wild-type (WT) and derivative strains deleted in *tssA* (Δ*tssA*), *vgrG* (Δ*vgrG*), the region encoding the VgrG glucosaminidase domain (*ΔvgrG*^*Tox*^), or the region spanning the VgrG glucosaminidase domain and the downstream *vgi* gene (*ΔvgrG*^*Tox*^*-vgi*) in LB at 28°C. Data are shown as the mean ± SD (n=4 technical replicates).

**Fig. S3**. Amino acid conservation of the VgrG glucosaminidase domain. A conserved motif found in the C-terminal region of VgrG (amino acids 700-820) is illustrated using WebLogo 3 based on multiple sequence alignment of proteins homologous to the VgrG glucosaminidase domain. A cyan circle above the WebLogo denotes E752, which is conserved in glucosaminidase domains and corresponds to an annotated active site residue in a member of the glucosaminidase family. Numbers below correspond to residue positions in VgrG.

**Fig. S4**. Expression of VgrG forms in *E. coli*. Myc-tagged VgrG wild-type (VgrG), VgrG point mutated at E752 (VgrG^E752A^), and core VgrG (amino acids 1-674; VgrG^Core^) were expressed in the cytoplasm or periplasm of *E. coli* from an arabinose-inducible plasmid, as indicated. Protein expression was detected by immunobloting using α-Myc antibodies. EV, empty vector.

**Fig. S5**. VgrG homologs in *Pantoea* carry diverse peptidoglycan-hydrolyzing toxin domains. Illustration of *Pab* VgrG homologs from other *Pantoea* strains representing the diversity of C-terminal toxin domains. The toxin domains in the top three examples were predicted by the NCBI Conserved Domain Database; the toxin domains in the bottom two examples were predicted by hidden Markov modeling (HHpred). PG, peptidoglycan.

**Fig. S6**. Pse2 sequence conservation and expression in *E. coli*. **A**. Amino acid conservation of Pse2 is illustrated using WebLogo 3 based on multiple sequence alignment of proteins homologous to Pse2. TMH, transmembrane helix. **B**. Myc-tagged Pse2 was expressed in the cytoplasm (cyto) or periplasm (peri) of *E. coli* from an arabinose-inducible plasmid. Protein expression was detected by immunoblotting using α-Myc antibodies. EV, empty vector.

**Movie S1**. Time-lapse microscopy of *E. coli* cells expressing periplasm-targeted VgrG. The GFP channel visualizing the fluorescent signal of the cell wall stain Wheat Germ Agglutinin Alexa Fluor 488 Conjugate (green) is shown. Cells were imaged every 5 min for 3 h.

**Movie S2**. Time-lapse microscopy of *E. coli* cells expressing periplasm-targeted VgrG containing a substitution in E752 to Ala. The GFP channel visualizing the fluorescent signal of the cell wall stain Wheat Germ Agglutinin Alexa Fluor 488 Conjugate (green) is shown. Cells were imaged every 5 min for 3 h.

**Movie S3**. Time-lapse microscopy of *E. coli* cells containing an empty expression plasmid. The GFP channel visualizing the fluorescent signal of the cell wall stain Wheat Germ Agglutinin Alexa Fluor 488 Conjugate (green) is shown. Cells were imaged every 5 min for 3 h.

