## Supporting Information Fig. S1 for "A dynamic antibacterial T6SS in *Pantoea agglomerans* pv. *betae* delivers a lysozyme-like effector to antagonize competitors"

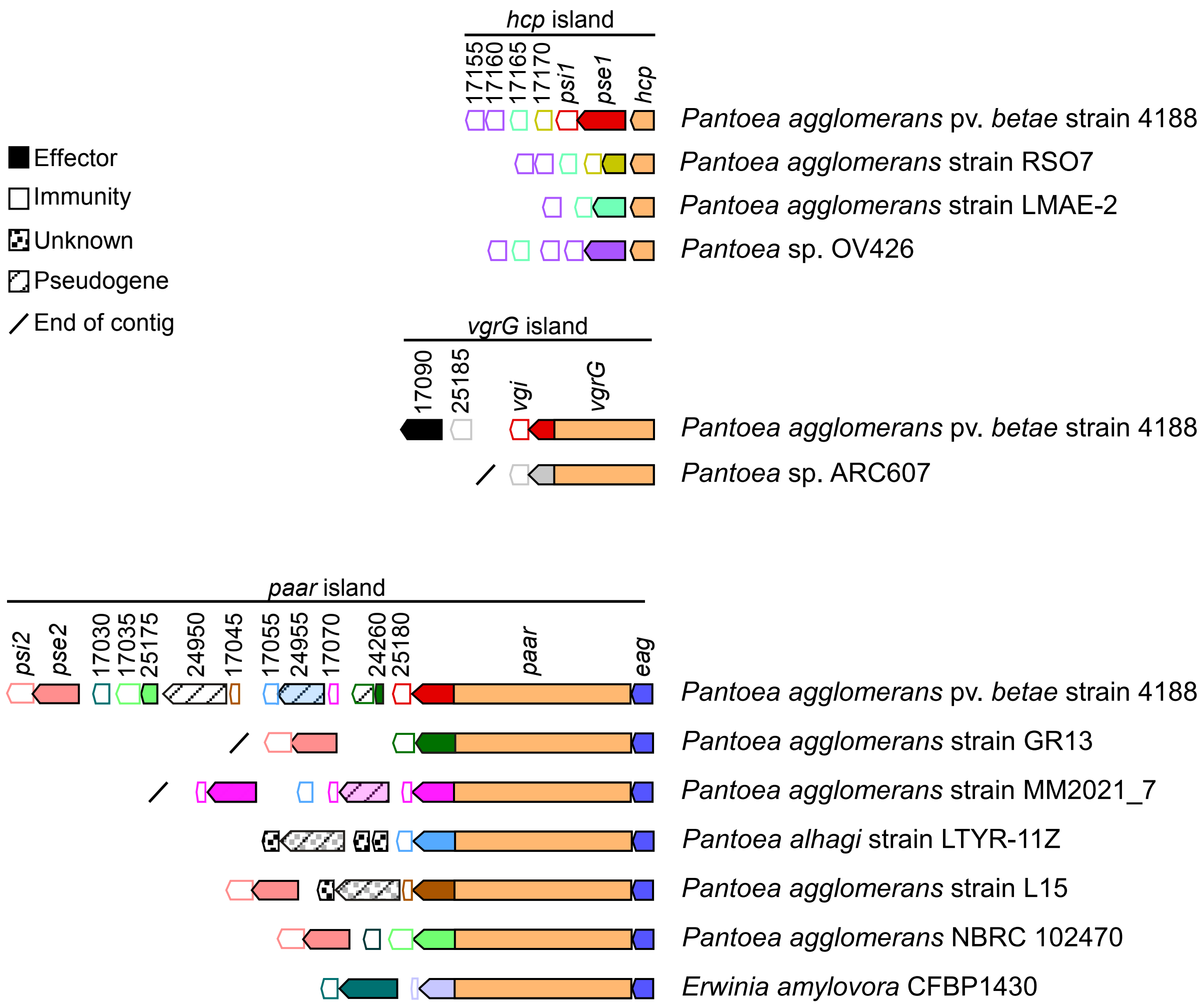


**Fig. S1.** Evolutionary dynamics in *Pab* T6SS1 islands. Genes are represented by arrows indicating the direction of transcription. Gene names or locus tags (A7P61_RSxxxxx) in the *Pab* T6SS1 cluster are denoted above. Colors denote homology between genes found in different genomes. Associated effector or toxin (full arrows) and immunity (empty arrows) pairs are denoted by the same color.
