## Supporting Information Fig. S2 for "A dynamic antibacterial T6SS in *Pantoea agglomerans* pv. *betae* delivers a lysozyme-like effector to antagonize competitors"

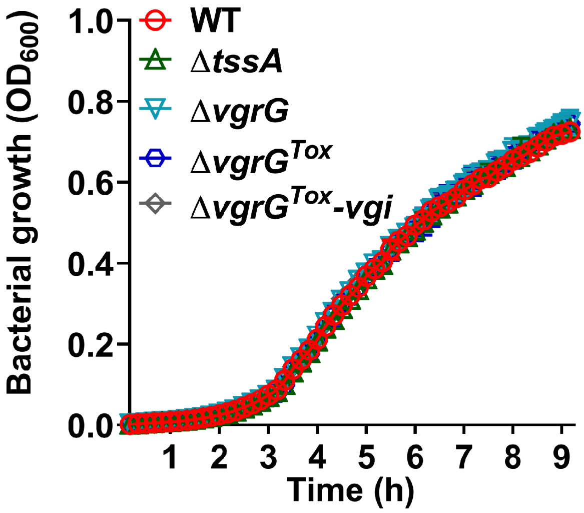


**Fig. S2.** *Pab* and derivative strains display a similar growth rate. Growth of *Pab* wild-type (WT) and derivative strains deleted in *tssA* (Δ*tssA*), *vgrG* (Δ*vgrG*), the region encoding the VgrG glucosaminidase domain (Δ*vgrG*$^{Tox}$), or the region spanning the VgrG glucosaminidase domain and the downstream *vgi* gene (Δ*vgrG*$^{Tox}$-*vgi*) in LB at 28°C. Data are shown as the mean ± SD (n=4 technical replicates).
