## Supporting Information Fig. S3 for "A dynamic antibacterial T6SS in *Pantoea agglomerans* pv. *betae* delivers a lysozyme-like effector to antagonize competitors"

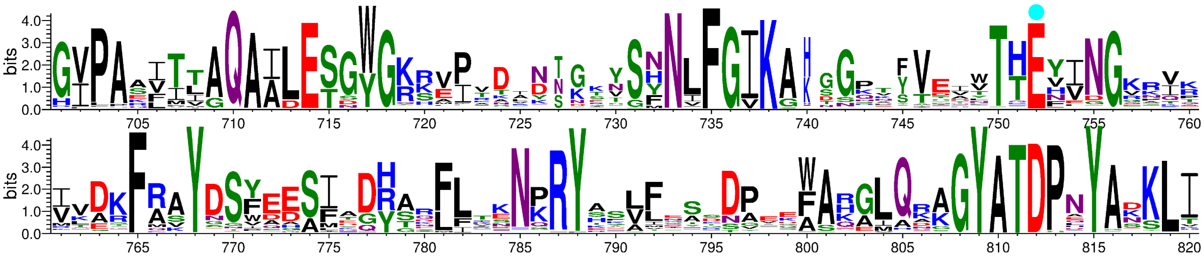


**Fig. S3.** Amino acid conservation of the VgrG glucosaminidase domain. A conserved motif found in the C-terminal region of VgrG (amino acids 700-820) is illustrated using WebLogo 3 based on multiple sequence alignment of proteins homologous to the VgrG glucosaminidase domain. A cyan circle above the WebLogo denotes Glu752, which is conserved in glucosaminidase domains and corresponds to an annotated active site residue in a member of the glucosaminidase family. Numbers below correspond to residue positions in VgrG.
