## Supporting Information Fig. S4 for "A dynamic antibacterial T6SS in *Pantoea agglomerans* pv. *betae* delivers a lysozyme-like effector to antagonize competitors"

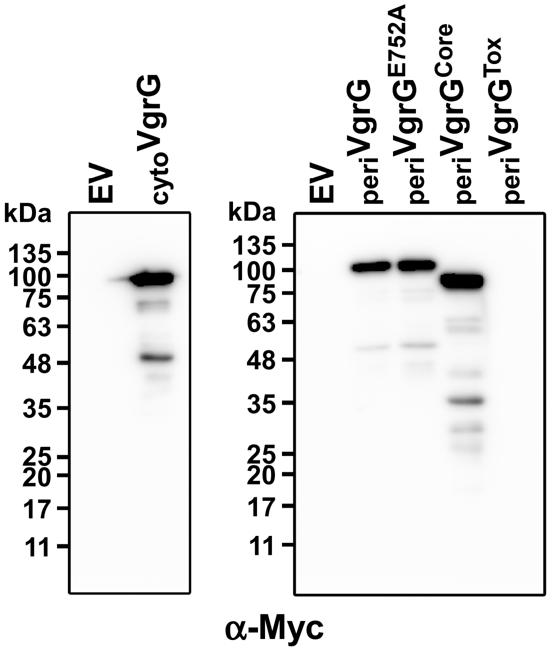


**Fig. S4.** Expression of VgrG forms in *E. coli*. Myc-tagged VgrG wild-type (VgrG), VgrG point mutated at E752 ($VgrG^{E752A}$), and core VgrG (amino acids 1-674; $VgrG^{Core}$) were expressed in the cytoplasm (cyto) or periplasm (peri) of *E. coli* from an arabinose-inducible plasmid, as indicated. Protein expression was detected by immunobloting using α-Myc antibodies. EV, empty vector.
