## Supporting Information Fig. S5 for "A dynamic antibacterial T6SS in *Pantoea agglomerans* pv. *betae* delivers a lysozyme-like effector to antagonize competitors"

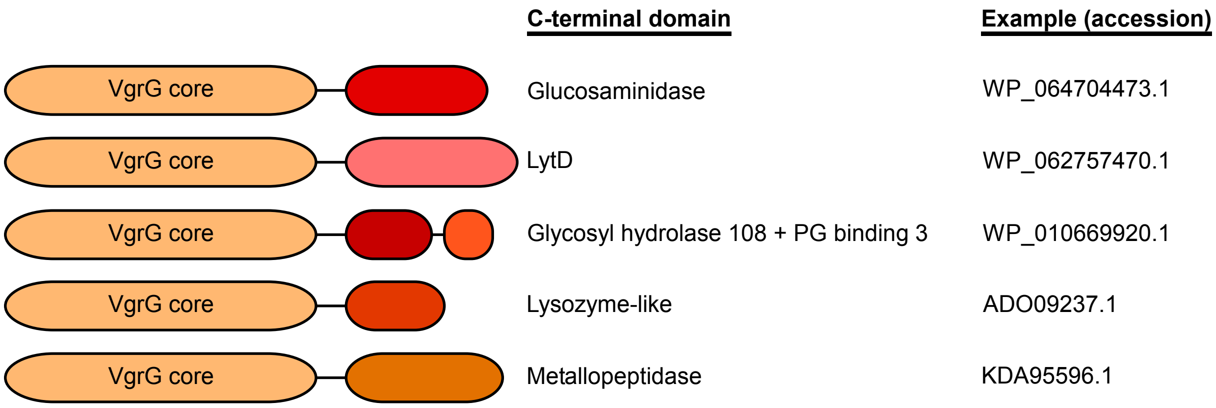


**Fig. S5.** VgrG paralogs in *Pantoea* carry diverse peptidoglycan-hydrolyzing toxin domains. Illustration of *Pab* VgrG paralogs from other *Pantoea* strains representing the diversity of C-terminal toxin domains. The toxin domains in the top three examples were predicted by the NCBI Conserved Domain Database; the toxin domains in the bottom two examples were predicted by hidden Markov modeling (HHpred). PG, peptidoglycan.
