## Supporting Information Fig. S6 for "A dynamic antibacterial T6SS in *Pantoea agglomerans* pv. *betae* delivers a lysozyme-like effector to antagonize competitors"

**A**

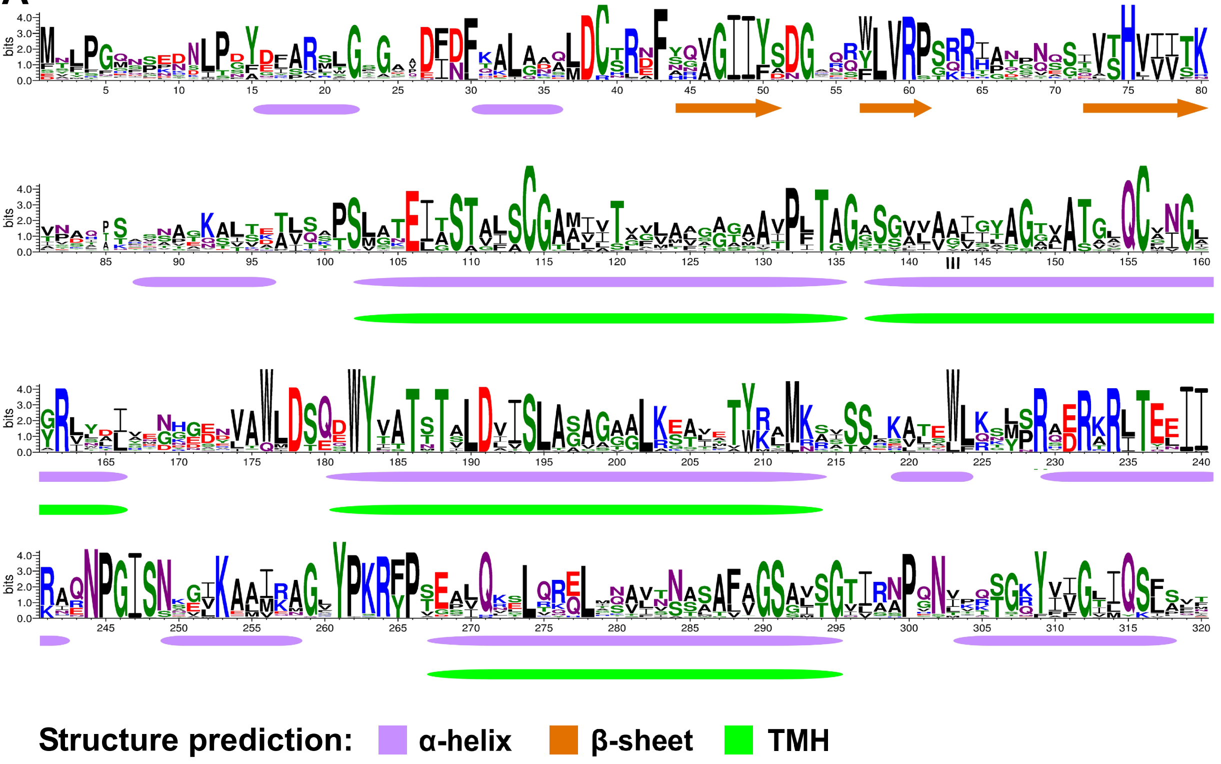


**B**

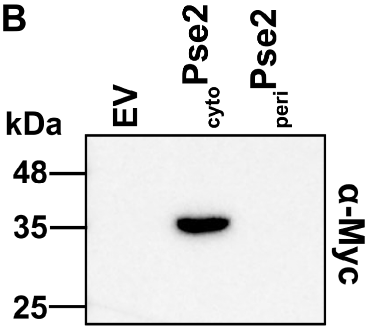


**Fig. S6.** Pse2 sequence conservation and expression in *E. coli*. **A.** Amino acid conservation of Pse2 is illustrated using WebLogo 3 based on multiple sequence alignment of proteins homologous to Pse2. TMH, transmembrane helix. **B.** Myc-tagged Pse2 was expressed in the cytoplasm (cyto) or periplasm (peri) of *E. coli* from an arabinose-inducible plasmid. Protein expression was detected by immunoblotting using α-Myc antibodies. EV, empty vector.
